# Phasic Activation of Ventral Tegmental, but not Substantia Nigra, Dopamine Neurons Promotes Model-Based Pavlovian Reward Learning

**DOI:** 10.1101/232678

**Authors:** R. Keiflin, H.J. Pribut, N.B. Shah, P.H. Janak

**Affiliations:** Department of Psychological and Brain Sciences, Krieger School of Arts and Sciences, Johns Hopkins University, Baltimore MD 21218 USA; The Solomon H. Snyder Department of Neuroscience, Johns Hopkins School of Medicine, Johns Hopkins University, Baltimore MD 21205 USA; Kavli Neuroscience Discovery Institute, Johns Hopkins University School of Medicine, Baltimore, MD 21205, USA

**Author notes:** Correspondence: Patricia H. Janak, Ph.D., Johns Hopkins University, 3400 N. Charles Street, Dunning Hall, room 246, Baltimore, MD 21218. Ronald Keiflin, Ph.D., Johns Hopkins University, 3400 N. Charles Street, Dunning Hall, room 246, Baltimore, MD 21218.

## Abstract

Dopamine (DA) neurons in the ventral tegmental area (VTA) and substantia nigra (SNc) encode reward prediction errors (RPEs) and are proposed to mediate error-driven learning. However the learning strategy engaged by DA-RPEs remains controversial. Model-free associations imbue cue/actions with pure value, independently of representations of their associated outcome. In contrast, model-based associations support detailed representation of anticipated outcomes. Here we show that although both VTA and SNc DA neuron activation reinforces instrumental responding, only VTA DA neuron activation during consumption of expected sucrose reward restores error-driven learning and promotes formation of a new cue→sucrose association. Critically, expression of VTA DA-dependent Pavlovian associations is abolished following sucrose devaluation, a signature of model-based learning. These findings reveal that activation of VTA-or SNc-DA neurons engages largely dissociable learning processes with VTA-DA neurons capable of participating in model-based predictive learning, while the role of SNc-DA neurons appears limited to reinforcement of instrumental responses.

## INTRODUCTION

The activity of midbrain dopamine (DA) neurons respond in a characteristic fashion to reward, with increased phasic firing in response to unexpected rewards or reward-predicting cues, little or no response to perfectly predicted rewards, and pauses in firing when predicted rewards fail to materialize^1–3^. This pattern of response largely complies with the concept of a signed reward prediction error (RPE), an error-correcting teaching signal featured in contemporary theories of associative learning^4–8^. Therefore, it has been suggested that the error signal carried by phasic DA responses and broadcast to forebrain regions constitutes the neural implementation of such theoretical teaching signals^5,9^. In support of this hypothesis, recent optogenetic studies showed that activation or inhibition of ventral tegmental area (VTA) DA neurons mimics positive or negative RPEs, respectively, and affects Pavlovian appetitive learning accordingly^10,11^. However, the specific learning strategy engaged by DA teaching signals remains controversial.

Computational models discriminate between two separate forms for error-driven learning^7,12^. In model-free learning, RPEs imbue predictive cues with a common currency cache value determined by the value of the outcome during training. This form of learning does not allow for a representation of the specific identity of the outcome; therefore, expression of this learning is independent of the desire for that specific outcome at the time of test. Alternatively, in model-based learning, error signals contribute to construction of internal models of the causal structure of the world, allowing predictive cues to signal the specific identity of their paired outcome. As a result, the expression of model-based learning is motivated by an internal representation of a specific outcome and anticipation of its inferred current value.

The role of DA teaching signals in model-free and model-based processes remains unclear^13–15^. Since the original discovery that DA neurons track changes in expected value, phasic dopamine signals have predominantly been interpreted as model-free RPEs, promoting pure value assignment. Consistent with this view, direct activation of DA neurons serves as a potent reinforcer of instrumental behavior in self-stimulation procedures^11,16–21^.

More recently however, a contribution of phasic DA signals to model-based learning has also been suggested. This is based on growing evidence that DA neurons have access to higher order knowledge, beyond observable stimuli, for the computation of RPEs^22–25^. Moreover, DA neurons were recently shown to respond to valueless changes in the sensory features of an expected reward^26^. While these studies reveal model-based influences in DA error signal computation, the exact associative content promoted by these DA signals is uncertain. A recent study intriguingly showed that in absence of a valuable outcome, phasic activation of DA neurons promotes model-based association between two neutral cues. Since the cues were neutral, there was no opportunity for model-free, value-based conditioning. It remains to be determined how DA signals contribute to associative learning when subjects are actively learning about value-laden rewarding outcomes, the canonical situation in which DA signals are robustly observed, and in which both model-free and model-based strategies are possible.

Potentially relevant to questions about the model-free or model-based nature of DA-induced learning is the proposed functional heterogeneity of DA neurons based on anatomical location. Indeed, while RPEs are relatively uniformly encoded across midbrain DA neurons, different contributions to learning have been proposed for VTA and substantia nigra (SNc) DA neurons based on the distinct ventral and dorsal striatal targets of these neurons^27–30^. Note however that unlike VTA-DA neurons for which a role in prediction learning is established, the contributions of phasic activity in SNc-DA neurons to error-driven prediction learning remains uncertain.

Therefore, the purpose of the present study was twofold: 1) assess the contribution of VTA- and SNc-DA neuron activation to Pavlovian reward learning, and 2) when learning was observed as a result of our manipulations, determine the model-free or model-based nature of this learning.

To accomplish these goals, rats were trained in a blocking paradigm in which the formation of an association between a target cue and a paired reward is prevented, or blocked, if this cue is presented simultaneously with another cue that already signals reward. In this situation, the absence of RPEs, presumably reflected in the absence of a DA response, is thought to prevent learning about the target cue. We sought to unblock learning by restoring RPEs, either endogenously by increasing the magnitude of reward, or by optogenetically activating VTA- or SNc-DA neurons during reward consumption. When successful unblocking was observed as a result of our manipulations, we assessed the model-free or model-based nature of the newly learned association by determining its sensitivity to post-conditioning outcome devaluation.

## RESULTS

### Phasic activation of VTA- but not SNc-DA neurons mimics reward prediction errors and promotes Pavlovian learning

Three groups of rats (Reward Upshift, n = 24; VTA-DA Stim, n = 20; SNc-DA Stim n = 16) were trained in a Pavlovian blocking/unblocking task (Fig. 1). In the first stage of this task, two auditory cues, A and B, were presented individually and followed by the delivery of a sucrose reward. For the Reward Upshift group, the quantity of sucrose associated with these cues was different: cue A signaled a large sucrose reward (3 × 0.1ml), while cue B signaled a small sucrose reward (0.1ml). This was done so that subsequent upshift of sucrose reward during the compound BY (from small to large reward) would cause an endogenous RPE and presumably unblock learning about the target cue Y. For the other two groups (VTA-DA Stim and SNc-DA Stim) cue A and B both signaled a large sucrose delivery, which, in absence of further manipulation should prevent endogenous RPEs during the subsequent compound phase. Subjects acquired conditioned responding rapidly, as indicated by the time spent in the reward port during cue presentation (Fig. 2). Note that during reinforced sessions, conditioned behavior was measured during the first 9s of cue presentation (before reward was delivered), in order to capture reward anticipation and avoid contamination by consumption behavior. In the reward upshift group, responding to cue A was greater than cue B (average for last 4 days of individual cue, T = 9.703, P < 0.001), which is consistent with the different reward magnitudes associated with these two cues. This difference in responding was not observed in VTA and SNc stim groups as in these groups, both cues signaled a large reward (Ps = 1.000; average last 4 days of individual cue).

**Figure 1.**
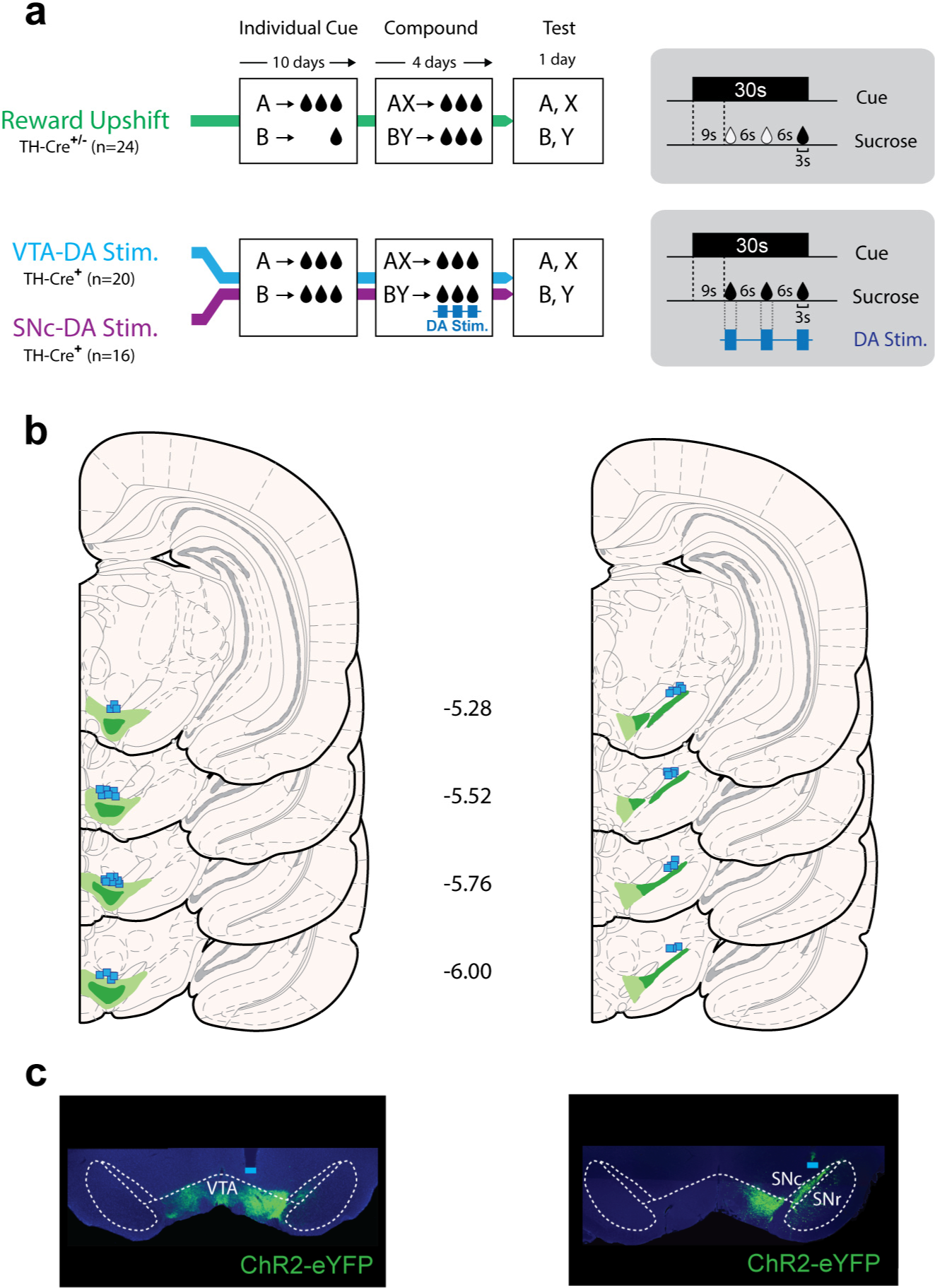
Behavioral task and histology. **a)** Three groups of rats were trained in the blocking/unblocking task. During the *Individual Cue* phase, two visual cues (A and B) were paired with sucrose reward. In the following *Compound Cue* phase, two auditory cues (X or Y) accompanied the initial visual cues to form two distinct compound stimuli (AX and BY). The absence of RPE during the compound AX is predicted to block learning about cue X. During the compound BY, a prediction error was produced either by increasing the magnitude of the sucrose reward (Reward Upshift group), or by photostimulating VTA- or SNc-DA neurons during sucrose consumption (VTA-DA Stim. and SNc-DA Stim. groups). A 1-day probe test assessed the associative strength acquired by each individual cue. **b)** Histological reconstruction of ChR2-YFP expression and fiber placement in VTA (left) and SNc (right). Light and dark shading indicate maximal and minimal spread of ChR2-YFP respectively. Square symbols mark ventral extremity of fiber implants. **c)** Representative ChR2-YFP expression after virus injection in VTA (left) or SNc (right).

**Figure 2.**
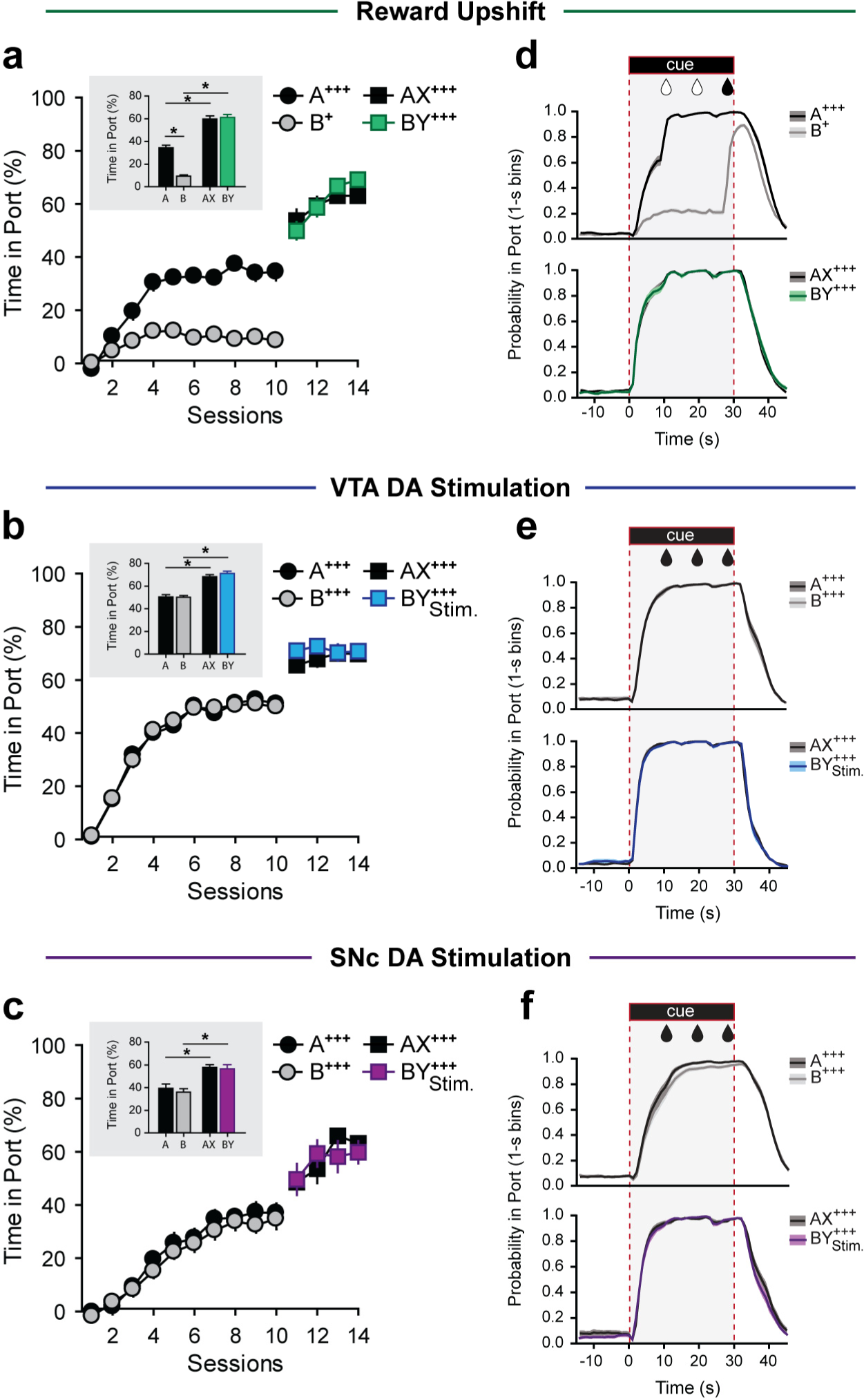
Performance during Individual Cue and Compound Cue training. **(a-c)** Time spent in reward port during cue presentation over 10 days of Individual Cue conditioning and 4 days of Compound Cue conditioning, for Reward Upshift **(a)**, VTA-DA stimulation **(b)**, and SNc-DA stimulation **(c)** groups. Values reported include only the first 9-s of the cues prior to sucrose delivery to avoid contamination with the consumption period. Inserts depict average performance over the last 4 days of Individual Cue conditioning (cue A and B) and over the 4 days of Compound Cue conditioning (compound AX and BY). For all groups, introduction of the auditory stimulus increased performance (A *vs*. AX, and B *vs*.BY, all Ps < 0.001, Bonferroni paired t-tests), but there was no difference in responding between the two compound cues (AX *vs*. BY, Ps > 0.967, Bonferroni paired t-tests). **(d-f)** Probability of presence in port throughout cue presentation during the last 4 days of Individual Cue (upper graphs) and in the 4 days of Compound Cue conditioning (lower graphs), for Reward Upshift **(d)**, VTA-DA stimulation **(e)**, and SNc-DA stimulation **(f)** groups. Note that photostimulation of VTA-DA or SNc-DA neurons during the compound cue, BY, did not disrupt ongoing behavior.

In the second stage of the procedure, two distinct visual cues (X and Y) accompanied the auditory cues to form to the compounds AX and BY. Both of these compound cues were paired with a large sucrose reward. For all subjects, the addition of cue X was redundant: a large reward was expected, and obtained, on the basis of cue A alone. Therefore, in absence of prediction error during AX trials, learning about the target cue X should be blocked. In contrast, the introduction of cue Y coincided with a prediction error. For the reward upshift group, the transition from small to large reward (from cue B to compound BY) is thought to create an endogenous prediction error that unblocks learning about the target cue Y. For the other two groups, we sought to artificially recreate a normally absent prediction error, by optogenetically activating VTA- or SNc-DA neurons during reward consumption in BY trials. For all groups, the transition from individual cue to compound trials was accompanied by a general increase in conditioned responding (A *vs*. AX, and B *vs*. BY Ps < 0.001) possibly reflecting both changes in associative weight and unconditioned arousing and/or disinhibiting properties of novel auditory cues^31^.

Finally, to assess the associative strength acquired by each individual cue following reward upshift or DA neuron optogenetic activation, all rats underwent a probe test in which all cues were presented separately and in absence of sucrose reinforcement (Fig. 3). Conditioned behavior was measured during the entire 30s cue. A two-way mixed ANOVA (Group × Cue) revealed a main effect of Group (F_2,57_ = 13.818, P < 0.01) and Cue (F_3,171_ = 17.997, P < 0.01) and a significant interaction between these factors (F_6,171_ = 11.050, P < 0.01). Follow-up one-way RM ANOVAs separately conducted on each experimental group revealed, for each group, a significant effect of cue type on responding (reward upshift: F_3,69_ = 22.078, P < 0.001; VTA-DA stimulation: F_3,57_ = 11.634, P < 0.001; SNc-DA stimulation: F_3,45_ = 7.836, P < 0.001). Posthoc comparisons confirmed that responding to the ancillary cues A and B was as expected: subjects in the reward upshift group responded more to A than to B (T = 5.373, P < 0.001), and subjects in the other two groups responded equally to these cues (VTA-DA stimulation: T = 0.904, P = 1.000, SNc-DA stimulation: T = 0.537, P = 1.000), which is consistent with the magnitude of reward paired with these cues during training. Of primary interest are the responses to the target cues X and Y. In the reward upshift group, the surprising increase in reward magnitude during the BY compound unblocked learning, resulting in greater conditioned responding to Y than to X (T = 5.841, P < 0.001). Note that both cues, Y and X, benefited from equal pairing with the sucrose reward during the compound phase, only the presence or absence of RPE during these cues differed and promoted or blocked learning, respectively. Stimulation of VTA-DA neurons during sucrose consumption in presence of the BY compound also resulted in greater responding to Y than to X (T = 5.334, P < 0.001). These results indicate that phasic activation of VTA-DA neurons mimicked endogenous RPEs and unblocked learning. In contrast, activation of SNc-DA neurons did not unblock Pavlovian learning; subjects responded equally to X and Y (T = 0.344, P = 1) and responding to these cues was low (< 10% of cue time spent in port, on any trial). Analysis of the rate of port entries during the cues (a different metric for the assessment of Pavlovian conditioned approach) yielded essentially similar results (Fig. S1). Note however that unlike the time in port, the rate of port entries did not follow a monotonic increase during Pavlovian training, making that last metric a somewhat ambiguous readout for changes in associative strength. Therefore, we chose to focus our primary analyses on time in port.

**Figure 3.**
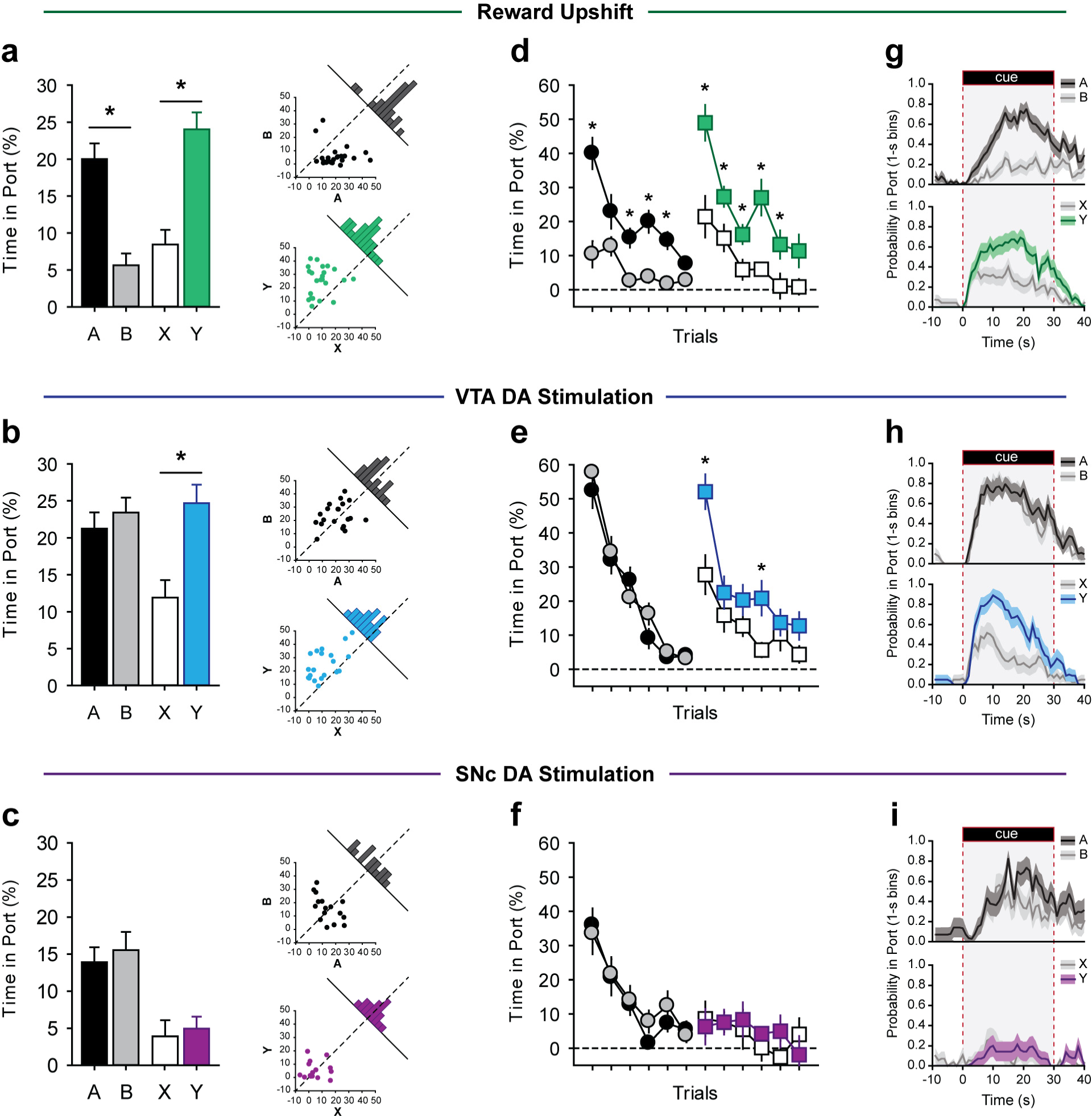
Photoactivation of VTA-DA but not SNc-DA neurons mimics endogenous RPEs and unblocks learning. Conditioned responding was measured by time spent in the reward port during cue presentation. **(a-c)**: Whole session performance in the Reward Upshift group **(a)**, VTA-DA stimulation group **(b)**, and SNc-DA stimulation group **(c)**. Scatterplot inserts on the right show individual data distributions for responding to cue A and B (top inserts), and X and Y (bottom insert). Histograms along the diagonal line are frequency distribution (subject counts) for the responding difference scores (A – B, or X – Y); off-centered distributions reveal higher responding to one of the cues. **(d-f)**. Trial-by-trial test performance in the reward upshift group **(d)**, VTA-DA stimulation group **(e)**, and SNc-DA stimulation group **(f)**. A 3-way mixed ANOVA (Group × Cue × Trial) analyzed the evolution of responding over the course of the session and found an interaction between all factors (F_30,855_ = 2.603, P < 0.001, after Greenhouse-Geisser correction). **(g-i)** Second-by-second tracking of presence in port during the first presentation of each cue (A and B: upper graph; X and Y: lower graph) for the reward upshift group **(g)**, VTA-DA stimulation group **(h)**, and SNc-DA stimulation group **(i)**. *P < 0.05 (A *vs*. B, or X *vs*. Y; Post-hoc Bonferroni t-test). Error bars = s.e.m.

To directly compare the consequence of endogenous RPEs and DA neurons activation on Pavlovian learning, we calculated for all individuals an unblocking score defined here as the difference in responding between Y and X (unblocked – blocked; using time in port as the measure of responding)(Fig. S2). We then compared this value between groups and while we found a general group effect (F_2,57_ = 8.247, P < 0.001), post hoc analysis found no difference between the reward upshift and VTA-DA stimulation groups (T = 0.817, P = 1) indicating equal unblocking as a result of these two manipulations. In contrast the unblocking score of the SNc-DA stimulation group was different from all other groups (all Ps ≤ 0.01), which confirms the functional dissociation between VTA- and SNc-DA neurons.

In certain conditions, cues paired with natural reward or with DA neurons stimulation can elicit behaviors that are not directed towards the reward port, such as orienting to the cue, rearing, and general locomotion/rotations^32,33^. To determine the role of endogenous- as well as optically induced-RPEs on the acquisition of these behaviors, we recorded and analyzed animals’ behavioral responses to cues X and Y during the probe test. While the target cues occasionally evoked orienting, rearing, or rotations, these behaviors were equally frequent in response to cue X and Y (Fig. S3), suggesting that, under these experimental parameters, these behaviors are not conditioned responses, but rather reflect the intrinsic (unconditioned) salient properties of the cues.

After completion of the unblocking task, we assessed the reinforcing properties of VTA- and SNc-DA neurons activation in an intracranial self-stimulation (ICSS) task in which rats could respond on one of two nosepokes to obtain a 1-s optical stimulation of DA neurons (Fig. 4). In agreement with previous studies^16,20,21,33^, we found that activation of both VTA- and SNc-DA neurons serves as a potent reinforcer of ICSS behavior. A 3-way mixed ANOVA (Group × Day × Nosepoke) conducted on responding over the course of two daily ICSS sessions revealed a clear preference for the active nosepoke (F_1,34_ = 45.522, P < 0.001) and a Nosepoke × Day interaction (F_1,34_ = 54.789, P < 0.001) as responding at the active nosepoke increased over time (T = 10.712, P < 0.001, Bonferroni *post hoc* tests) while responding at the inactive nosepoke remained virtually absent (T = 0.0414, P < 0.967, Bonferroni *post hoc* tests). Critically, we found no main effect (F_1,34_ = 0.876, P = 0.356) or interaction with group (Group × Day: F_1,34_ = 0.244, P = 0.625; Group × Nosepoke: F_1,34_ = 0.777, P = 0.384; Group × Day × Nosepoke: F_1,34_ = 0.270, P = 0.607), indicating that stimulation of VTA- and SNc-DA neurons is equally reinforcing in the ICSS paradigm.

**Figure 4.**
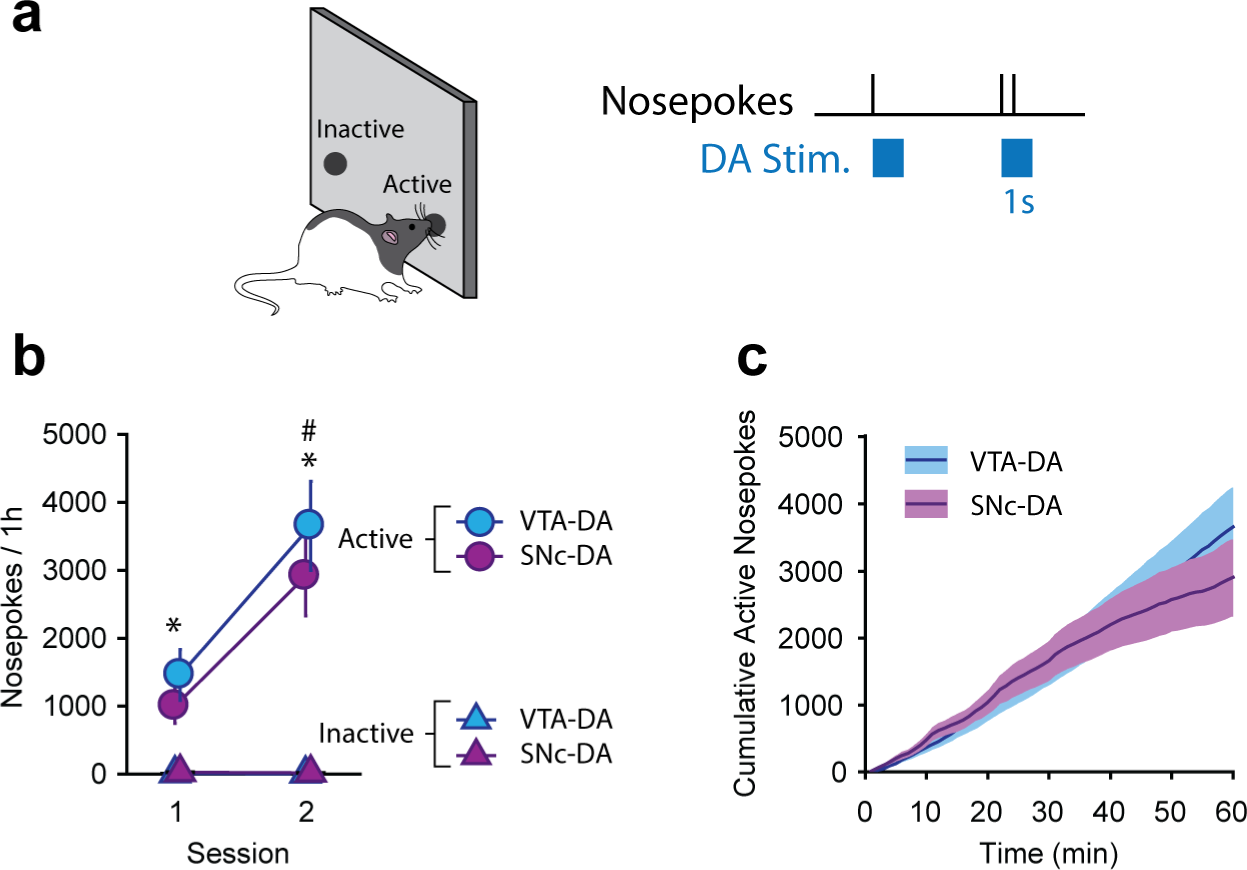
Photoactivation of VTA-DA or SNc-DA neurons serves as an equally potent reinforcer of ICSS behavior. **(a)** Rats could respond on one of two nosepokes to obtain an optical stimulation of VTA- or SNc-DA neurons. **(b)** Responses at the active and inactive nosepoke over the course of two daily 1-h sessions. **(c)** Cumulative responses at the active nosepoke over the course of the last ICSS session. *P < 0.05, Active vs. Inactive Nosepoke; #P < 0.05, Session 1 vs. Session 2 (active nosepoke). Error bar and error bands = s.e.m.

Together, these results show that while VTA- and SNc-DA neuron activation are equally potent reinforcers of instrumental behavior, only VTA-DA neurons activation mimics endogenous RPEs in promoting error-correcting Pavlovian learning (unblocking).

### Activation of VTA-DA neurons engages model-based learning

Although we demonstrated above that endogenous RPEs induced by reward upshift and optogenetic activation of VTA-DA neurons result in numerically comparable unblocking effects, the learning strategy engaged by these two manipulations might be different. In model-free algorithms, RPEs imbue predictive cues with a scalar cache value, resulting in conditioned responses largely independent of the outcome value at the time of test. Alternatively, in model-based accounts, predictive cues come to signal the specific identity of their paired outcome, resulting in conditioned responses motivated by the sensorily-rich representation of the outcome and its current value. To determine the learning strategy recruited by endogenous RPEs, or VTA-DA neuronal activation, we assessed the effect of devaluing the sucrose outcome on responding to Y, the unblocked cue. New groups of rats were trained in the blocking/unblocking task previously described: learning about cue Y was unblocked either by reward upshift (n = 24) or by VTA-DA neurons stimulation (n = 23) during the BY compound. At the end of the compound training phase, rats in each group were assigned to one of two conditions. Subjects in the “devalued” condition had sucrose devalued by pairing its consumption in the homecage with LiCl-induced nausea (conditioned taste aversion). For the subjects in the “valued” condition, sucrose consumption and LiCl-induced nausea occurred on alternate days, which preserved the value of the sucrose outcome (Fig. 5, Fig. S4). Two days after the final LiCl injection, all rats were then tested for conditioned responding to Y (unblocked cue) and to A (ancillary cue paired with large reward) in separate probe sessions with the order of testing counterbalanced.

**Figure 5.**
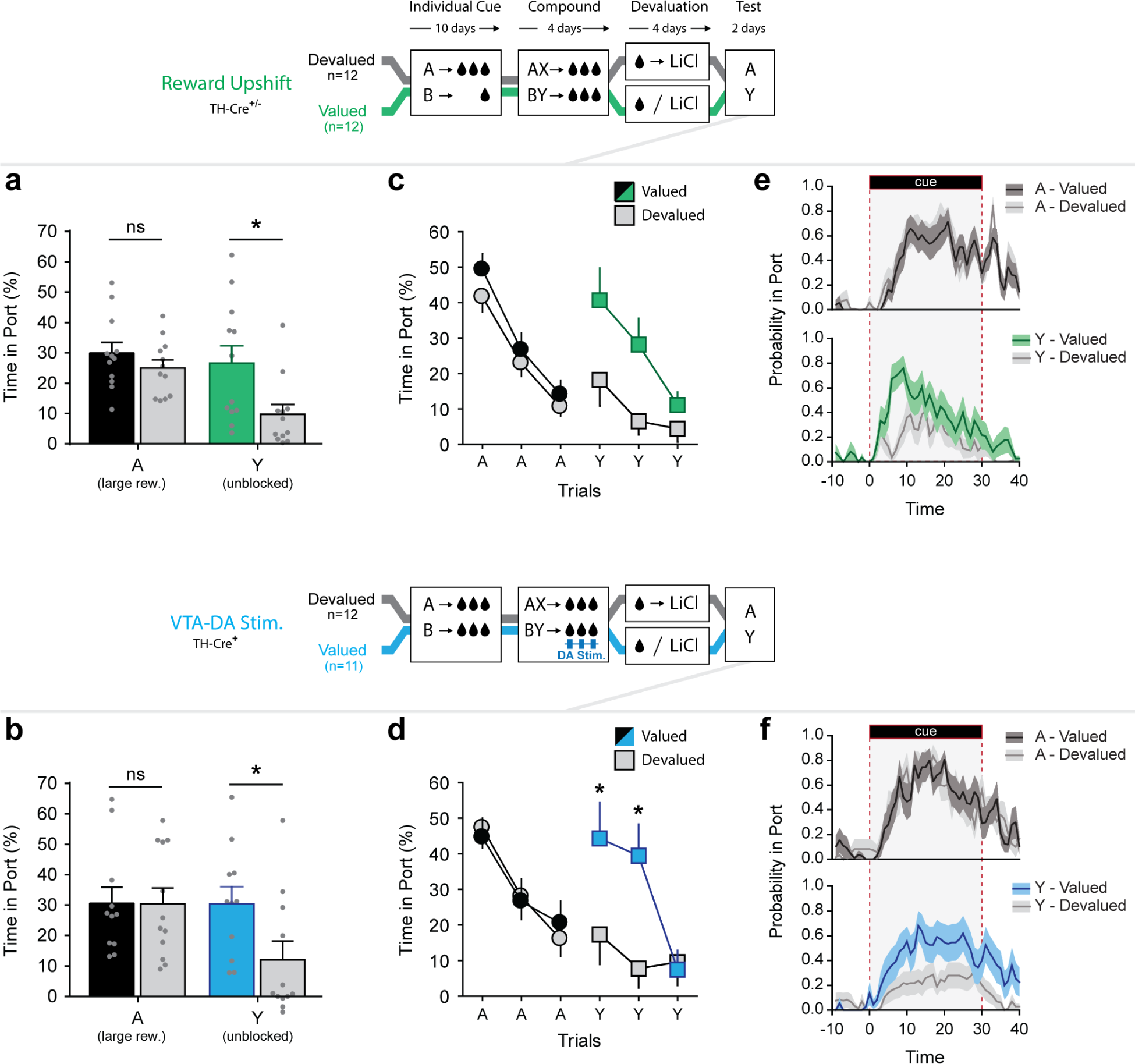
Devaluation of the sucrose outcome abolishes conditioned responding to the unblocked cue Y in Reward Upshift and VTA-DA groups. Learning about the target cue Y was unblocked either by reward upshift (top graphs), or by activation of VTA-DA neurons (bottom graphs). Following unblocking, the sucrose outcome was devalued for half the subjects in Reward Upshift and VTA-DA groups by pairing sucrose consumption with LiCl (Devalued condition). The remaining subjects were exposed to sucrose or LiCl-induced illness on alternate days preserving the value of sucrose (Valued condition). Conditioned responding to Y (unblocked cue) and A (cue paired with large reward) was then assessed at Test. **(a, b)** Time spent in reward port during cue presentation, in the Reward Upshift group **(a)** and VTA-DA group **(b)**. Sucrose devaluation reduced responding to Y in both groups. **(c, d)** Trial-by-trial performance in the Reward Upshift group **(c)** and VTA-DA stimulation group **(d)**. Independent 3-way ANOVAs (Cue × Devaluation × Trial) found an interaction between all these factors for the VTA-DA group (F_2,20_ = 3.901, P = 0.037), but not the Reward Upshift Group (F_2,21_ = 1.276, P = 0.300). **(e, f)** Second-by-second tracking of presence in port during the first presentation of each cue. *P < 0.05 (Valued vs. Devalued; Bonferroni t-test). Error bar and error bands = s.e.m.

A 3-way mixed ANOVA (Group × Devaluation × Cue) conducted on the time in port during the cues revealed a main effect of Cue (F_1,43_ = 6.119, P = 0.017) and of Devaluation (F_1,43_ = 10.707, P = 0.002) as well as an interaction between these factors (F_1,43_ = 4.750, P = 0.035). This interaction was due to a significant influence of the devaluation procedure on responding to the unblocked cue, Y (T = 3.563, P<0.001), but not on the ancillary cue, A (T = 0.514, P = 0.609). Reduced responding to Y after sucrose devaluation indicates that this response is normally motivated by the representation of the sucrose outcome and anticipation of its current value (model-based process). Critically, we found no main effect (F_1,43_ = 0.869, P = 0.356) or interaction with Group (Group × Devaluation: F_1,43_ = 0.005, P = 0.943; Group × Cue: F_1,43_ = 0.000, P = 0.993; Group × Devaluation × Cue: F_1,43_ = 0.339, P = 0.564). Planned contrast analyses independently confirmed that, for each group, sucrose devaluation reduced responding to unblocked cue Y (Reward Upshift: T= 2.559, P = 0.018; VTA-DA Stim.: T= 2.116, P = 0.046), but not to A (Reward Upshift: T= 1.126, P = 0.272; VTA-DA Stim.: T= 0.018, P = 0.986). Analysis of the rate of port entries during the cues yielded essentially similar results (Fig. S5). Note that VTA-DA valued and devalued subjects later displayed similar ICSS behavior (Fig. S6), which indicates that the reduced responding to the unblocked cue Y in devalued subjects cannot be explained by reduced efficiency of the optical stimulation, and thus reduced unblocking, in those animals. These results indicate that both endogenous RPEs and VTA-DA neuronal activation during sucrose consumption promoted the formation of model-based associations and conferred cue Y with the ability to evoke a representation of the sucrose outcome.

## DISSCUSION

We have shown that activation of VTA, but not SNc, DA neurons mimics RPEs and promotes the formation of model-based cue-reward associations. We used a Pavlovian blocking procedure, in which the formation of a cue-reward association is normally blocked by the absence of RPE (the reward being signaled by other predictive stimuli in the environment). Confirming and extending a previous study in our lab^11^, we showed that restoring RPEs, either endogenously, by increasing the magnitude of the sucrose reward, or by optogenetic activation of VTA-DA neurons, unblocks learning and promote the formation of a cue-reward association. In a separate experiment, we probed the content of this newly formed association by assessing its sensitivity to outcome devaluation. We found that following unblocking by reward upshift, or by VTA-DA stimulation, the expression of the unblocked learning was sensitive to the current value of the outcome; post-unblocking devaluation of the sucrose outcome almost entirely abolished responding to the unblocked cue. This indicates that both manipulations (reward upshift or VTA-DA stimulation), promote the formation of model-based associations that integrate a representation of the specific identity of the rewarding outcome. In stark contrast, we showed that optogenetic activation of SNc-DA neurons failed to promote Pavlovian learning, i.e., learning remained blocked. This is despite the fact that activation of both VTA- and SNc-DA neurons serves as a potent reinforcer in self-stimulation procedures.

These results are consistent with a recent study by Sharpe and colleagues showing that phasic VTA-DA responses mediate the formation of the association between two neutral stimuli (A→B), a form of learning that is necessarily model-based since it involves only identity and not value^34^. The status of this association was then assessed by pairing one of the stimuli with a food reward (B→food) and testing the conditioned responding to the other stimulus (A); food-seeking responses evoked by the target cue revealed a learned association between the two stimuli and inference of upcoming food reward (i.e., if A→B and B→food, then A→food). While Sharpe et al. demonstrated for the first time that VTA-DA signals can promote the association between neutral stimuli, this study did not address the nature of *reward* encoding in DA-dependent associations. Indeed, although their study involved a natural reward, it was used simply as a necessary means to reveal stimulus-stimulus associations, and was not the object of DA manipulations. This distinction is important because unlike stimulus-stimulus associations that are by definition model-based, cue-reward associations can be encoded in a model-free *or* model-based manner. Therefore, the possibility remains that while capable of promoting model-based learning when only sensory information is available, VTA DA signals nevertheless engage preferentially modelfree learning when (model-free) value can be encoded. In the present study, optogenetic activation of DA neurons was used to promote direct cue-reward associations, a form of learning that presents the opportunity for model-free and model-based algorithms. In these conditions, in which both learning strategies are equally valid, we showed that VTA-DA signals engage preferentially model-based learning.

Note however that our results do not preclude the participation of VTA-DA signals in model-free value assignment. Indeed, as shown here (ICSS experiment) and elsewhere^16,33^, in absence of external reward, the activation of VTA-DA neurons can confer cues and action with incentive/action value. Ultimately and consistent with DA’s neuromodulatory role, the content of DA-induced learning is likely dependent on the nature of the information being encoded and processed in terminal regions when coincident DA surges occur. What we show here is that in the presence of an external reward, the recruitment of a model-based learning strategy is not an exception but rather a central feature of VTA-DA teaching signals. This is consistent with recent studies showing that treatments (pharmacological or dietary restrictions) that globally increase or decrease DA function promote or impair, respectively, model-based processes in humans^35–37^. Note however that these treatments also affect tonic DA levels, which could affect learning independently of the phasic error signals.

An intriguing aspect of our results is the dissociation between the unblocked cue Y and the ancillary cue A in terms of response strategy. Indeed, unlike cue Y, cue A evoked conditioned responding that was driven by model-free associations (not affected by sucrose devaluation). The reason for this dissociation is unknown, but might involve training history differences between these cues. Indeed, compared to cue Y, cue A benefited from an extensive training history (224 conditioning trials vs. 32 for cue Y) which has been shown to promote model-free learning, although generally in the context of instrumental conditioning^38–40^. Perhaps more interesting are the implications for the role of VTA-DA signals in learning. In the VTA-DA group, the cues A and B are equivalent up to the compound conditioning phase and, based on the lack of effect of devaluation on A, we can assume that responding to both cues is governed by model-free associations. Therefore it appears that the activation of VTA-DA neurons promoted the formation of model-based associations about Y in subjects that were (presumably) currently engaged in model-free behavior during BY trials. This surprising result suggests that model-free associations could be formed “in the background”, independently of the strategy that governs behavior at the time these associations are formed, or through post-training event replay^41^. Alternatively, it could be that activation of VTA-DA neurons is sufficient to shift response strategy and restore model-based processing^42^. Further studies are required to address these questions.

Our results provide strong evidence for a functional dissociation between VTA- and SNc-DA neurons in appetitive learning. While activation of VTA-DA neurons unblocked Pavlovian learning, we found no evidence of unblocking following SNc-DA neurons activation, despite careful analysis of several behavioral responses (time in port, port entries, orienting, rearing, and locomotion). This contrasts with recent results from our lab showing that, in absence of a natural reward, activation of VTA- or SNc-DA neurons during cue presentation promotes the development of conditioned cue-evoked locomotion^33^. An important point to consider when comparing these results is the behavior of the animals at the time of the stimulation. Although free movement was possible, animals in the present study were relatively immobile during DA stimulation because it occurred as they were consuming the sucrose reward. This absence of ambulatory movement during DA stimulation could have prevented the emergence of conditioned locomotion.

In contrast with the selective role of VTA-DA neurons in Pavlovian unblocking, we show in here, in agreement with previous studies^20,33^, that instrumental behavior for ICSS can be supported by either VTA- or SNc-DA neurons stimulation. This partial dissociation between VTA- and SNc-DA neurons in Pavlovian and instrumental learning is reminiscent of the actor-critic reinforcement algorithm. This model is based on the idea of a separation of labor between a prediction module and an action module, with distributed RPEs promoting learning in both modules but with different consequences (updating predictions vs. reinforcing actions). A possible neural implementation of actor-critic algorithm has been suggested, with ventromedial (VMS) and dorsolateral (DLS) striatum functioning as prediction and action modules, respectively^29^. Consistent with this, we showed that activation of SNc-DA neurons, projecting predominantly to DLS, reinforces prior actions but has no influence on Pavlovian prediction learning, in agreement with the role of RPEs in an action module, while activation of VTA-DA neurons, projecting predominantly to VMS, promote Pavlovian learning, in agreement with the role of RPEs in a prediction module. Because predictions are updated by RPEs but also influence RPEs computations in return, the actor-critic model predicts that RPEs in the prediction module reinforce Pavlovian cues/states, which can then subsequently evoke back-propagated RPEs, including in the action module. A neural equivalent of this process, in which Pavlovian predictions encoded in the VMS feed back onto midbrain DA neurons (including SNc-DA neurons) and contribute to the propagation of a RPE teaching signal to more dorsal and lateral portions of the striatum, could contribute to the instrumental reinforcement induced by VTA-DA stimulation. However, a critical difference between our results and the predictions of the actor-critic algorithm is that this algorithm is strictly model-free, while we have shown here that VTA-DA signals contribute to model-based Pavlovian learning. Therefore, our results suggest a hybrid model that incorporates both model-free and model-based processes and in which VTA DA dependent model-based predictions shape SNc-DA signals and train model-free instrumental learning^43^

Finally, these results have important implications for our understanding of DA-related pathologies. Noisy/deregulated DA signals originating from the VTA, such has been observed in schizophrenic patients^44,45^, could promote model-based associations between external and/or internal events that are merely coincident but not causally-related, leading to the construction of internal world models that are out of touch with the physical reality and sources of delusional beliefs^46^. In contrast, emergence of cue- or reward-evoked DA signals in the DLS, such has been reported after repeated drug use^47–50^, could contribute to the reinforcement of maladaptive drug-seeking responses that persist despite knowledge of their adverse consequences^51,52^.

## ACKNOWLEDGEMENTS

This work was supported by National Institutes of Health grant DA035943.

## AUTHOR CONTRIBUTIONS

R.K. and P.H.J. conceived the study; R.K., H.J.P, and N.B.S. carried out the experiments; R.K. analyzed the data; R.K. and P.H.J. wrote the manuscript with input from all the authors.

### Author Information

The authors declare no competing financial interests. Correspondence and requests for materials should be addressed to RK or PHJ

## METHODS

### Subjects

*Th::Cre*+ transgenic rats (37 males and 24 females; Long-Evans background) expressing Cre recombinase under the control of the tyrosine hydroxylase promoter and their wild-type littermates (30 males and 16 females;*Th::cre*^−^) were used in these studies. Rats were singly housed under a 12 h light/12 h dark cycle with unlimited access to food and water, except during the behavioral experiment, when they were mildly food restricted to ~90% of their free-feeding weight. Behavioral experiments were conducted during the light cycle. All experimental procedures were conducted in accordance with the UCSF and JHU Institutional Animal Care and Use Committees and the US National Institute of Health guidelines. Males and females were distributed as evenly as possible across the different experimental groups. No significant effects of sex were found; therefore data for males and females were collapsed.

### Surgeries

Under stereotaxic guidance, anesthetized *Th::Cre*+ rats (>300g males; >225g females) received unilateral infusions of AAV5-EF1α-DIO-ChR2-eYFP (titer: 1.5−4×10^12^ virus particles/mL) into the VTA (AP: −5.4 and −6.2mm from bregma; ML: ± 0.7 from midline; DV: −8.5 and −7.5 from skull) or the SNc (AP: −5.0 and −5.8; ML: ± 2.4; DV: −8.0 and −7.0). This resulted in 4 injection sites for each rat. At each injection site, 1μl of virus was injected at the rate of 0.1μL/min. In the same surgery, the rats were also implanted with optic fibers aimed at the VTA (AP: −5.8; ML: ± 0.7; DV: −7.5) or the SNc (AP: −5.4; ML: ± 2.4; DV: −7.2). The optic fiber implants were made in-house and had a custom lock-in mechanism that ensured secure and durable connection to the patch cable during behavioral sessions. Behavioral experiments started >2 weeks post-surgery; sessions that included optical stimulation were conducted >4 weeks post-surgery.

### Apparatus

Behavioral sessions were conducted in 12 identical sound-attenuated conditioning chambers (Med Associates, St. Albans, VT). A liquid delivery port was recessed in the center of the right wall ~2 cm above the floor and connected to a syringe pump located outside of the sound-attenuating cubicle. The left wall had two nosepoke operanda (18 cm apart). A houselight was centered above the left wall and a pair of cue lights flanked the liquid delivery port on the right wall. Additionally, a white noise (76dB) and two pure tones (2.9 and 4.5 kHz, both 76dB) could be delivered through 3 wall speakers. The nosepoke operanda were obstructed during the unblocking procedures and accessible only during ICSS sessions. Conversely, the sucrose port was accessible only during the unblocking procedures but obstructed during ICSS sessions. Subjects’ presence in the port or nosepokes was detected by interruption of infrared beams.

### Unblocking by reward upshift

In a brief shaping session, rats were trained to consume sucrose (15%, w/v) delivered in the liquid port (0.1 ml/delivery; 30 deliveries over the course of 45 min). All rats then received 10 daily conditioning sessions during which two 30-s visual cues, A and B (the flashing of the houselight 1 s on, 2 s off, or the steady illumination of the light cues; counterbalanced) were paired with two different quantities of sucrose. Cue A signaled a large sucrose reward: 0.3 ml with 0.1 ml delivered every 9 s of the 30-s cue. Cue B signaled a small sucrose reward: 0.1 ml delivered over the last 3 s of the 30-s cue. These conditioning sessions consisted of 16 presentations of each cue with an average intertrial interval (ITI) of 3 min ± 1.5 (rectangular distribution; average ITI maintained constant throughout the experiment). After this initial phase of individual-cue conditioning, rats were pre-exposed to two auditory stimuli, X and Y (the beeping of the tones 0.1 s on, 0.2 s off, or a steady white noise; counterbalanced) in a single habituation session (six 30-s presentation of each cue, no sucrose delivered). Over the next 4 days, rats received conditioning to the compound cues. Simultaneous presentations of A and X (AX compound), or B and Y (BY compound) were paired with the large sucrose reward. Cues A and B also continued to be presented individually with their respective rewards as in training, as a reminder of the individual value of these cues. Each compound conditioning session consisted of 8 presentations of each trial type (AX, BY, A, B). Following compound conditioning, all rats received a probe test consisting of six unrewarded 30-s presentation of A, B, X, Y (in blocks of 3; order counterbalanced).

### Unblocking by VTA- or SNC-DA Stimulation

Behavioral procedures were similar to those described for the unblocking by reward upshift, with the following exceptions: 1) during initial conditioning to the individual cues, both cue A and B were paired with a large sucrose reward, which, in absence of further manipulation, should result in the blocking of both cue X and Y; and, 2) during compound conditioning, each delivery of sucrose during the compound BY was accompanied by a 3-s train of light pulses (473 nm, 20 Hz, 60 pulses, 5 ms duration) delivered into the VTA or the SNc. The delivery of a train of stimulation required 100 ms of continuous presence in the baited port, in order to coincide with the consumption of the sucrose reward. Rats were tethered to the optical patch cord for most conditioning sessions with the exception of training day 1, 5, 8, and pre-exposure to X and Y. This was done in order to habituate rats to perform the task in both conditions (tethered and untethered). For the final probe test, rats were not tethered to the optical patch cable, in order to prevent any potential interference of tethering on behavior (particularly, on orienting responses).

### Outcome devaluation

Rats were initially trained in the unblocking task where learning about the target cue Y was unblocked by reward upshift or by photoactivation of VTA-DA neurons. At the end of compound conditioning and before the final probe test, half of the rats in each group had the sucrose outcome devalued by pairing it with lithium chloride (LiCl)-induced nausea (devalued condition). Devaluation took place in the animals’ homecage over 4 days. On day 1 and 3, rats in the devalued groups received 10 min of free access to sucrose immediately followed by an injection of LiCl (0.3 M; 6 ml/kg). The other half of the rats (valued condition) received similar exposure to sucrose and LiCl-induced illness but on alternate days (LiCl injections on Day 1 and 3; sucrose access on day 2 and 4). To confirm that sucrose devaluation was durable and transferred across contexts, sucrose consumption was then measured in the conditioning chambers. Rats were placed in the chambers for 5 min, where the reward cup had been filled with 4 ml of sucrose. After 5 min, rats returned to their homecage and the remaining amount of sucrose was measured to calculate consumption. This brief sucrose consumption test occurred twice, a day before and a day after the cue probe tests. No difference was found between these two consumption tests, therefore the results were collapsed. Cue probe tests consisted of 6 unrewarded presentation of Y (unblocked cue) and A (control cue of comparable high value), on alternate days (order counterbalanced) in order to prevent potential interference between different response strategies (model-free vs. model-based). In these conditions, conditioned responding rapidly extinguished within session, therefore only responding on the first 3 trials was analyzed.

### Intra-Cranial Self-Stimulation (ICSS)

Following completion of the unblocking procedures, all Th::Cre^+^ rats in the VTA- and SNc-DA groups were tested for intra-cranial self-stimulation. During two daily 1-h sessions, rats had access to two nosepoke ports; a response at the active nosepoke (position counterbalanced) resulted in the delivery of a 1-s train of light pulses (20 Hz, 5 ms duration) into the VTA or the SNc. Responses at the active nosepoke during the 1-s light train were recorded but had no consequence. Responses at the inactive port were always without consequence.

### Video Analysis

A camera located in each conditioning chamber and connected to a video acquisition software (Noldus Information Technology, Leesburg, VA) recorded animals’ behavior during the probe test session. The purpose of these recordings was to capture behavioral responses that are not directed towards the reward port (and therefore cannot be automatically detected by infrared beam interruption). Three types of responses were detected and manually scored: i) *orienting responses*, defined as rapid head movements in the direction of the cue, which occurred within 3s of the cue onset. ii) *rearing responses*, defined as the animal standing on hind legs with front feet off the floor (often against the side walls) and not grooming. iii) *rotation responses*, defined as the animal making a full rotation in the chamber between the onset and termination of the cue.

### Histology

Implanted animals were deeply anesthetized with pentobarbital and perfused transcardially with 0.9% saline followed by 4% paraformaldehyde. Brains were extracted, cryoprotected in 25% sucrose for >48 hours, and sectioned at 50 μm on a freezing microtome. Coronal slices were collected onto glass slides and coverslipped with Vectashield mounting medium with DAPI. Fiber tip position and eYFP-CHR_2_ virus expression were examined under a fluorescence microscope (Zeiss Microscopy, Thornwood, NY).

### Data Analysis

Counterbalancing procedures were used to form experimental groups balanced in terms of sex, cue identity, and behavioral performance in the sessions preceding the experimental intervention. Conditioned responding was measured primarily by the percentage of time in the port, or the rate of port entries, during cue presentation, normalized by subtracting behavior during a pre-cue period of equal length. Behavior during pre-cue period was always extremely low (0.304s ± 0.057 of average presence in the port during the 30s that precede cue presentation, no group difference Ps > 0.752). Statistical analyses were conducted using SPSS Statistics V22, and Systat SigmaPlot 14, and consisted generally of mixed-design repeated measures ANOVAs with cue and trials as within-subject factors, and group (reward upshift, VTA-DA, or SNc-DA) and devaluation as between-subject factors. On the rare occasions that the sphericity assumption was violated, the Greenhouse-Geisser correction was used to adjust the reported p-value. Post-hoc and planned comparisons were carried with Bonferroni t-test. Significance was assessed against a type I error rate of 0.05.

## SUPPLEMENTAL MATERIAL FOR

**Figure S1.**
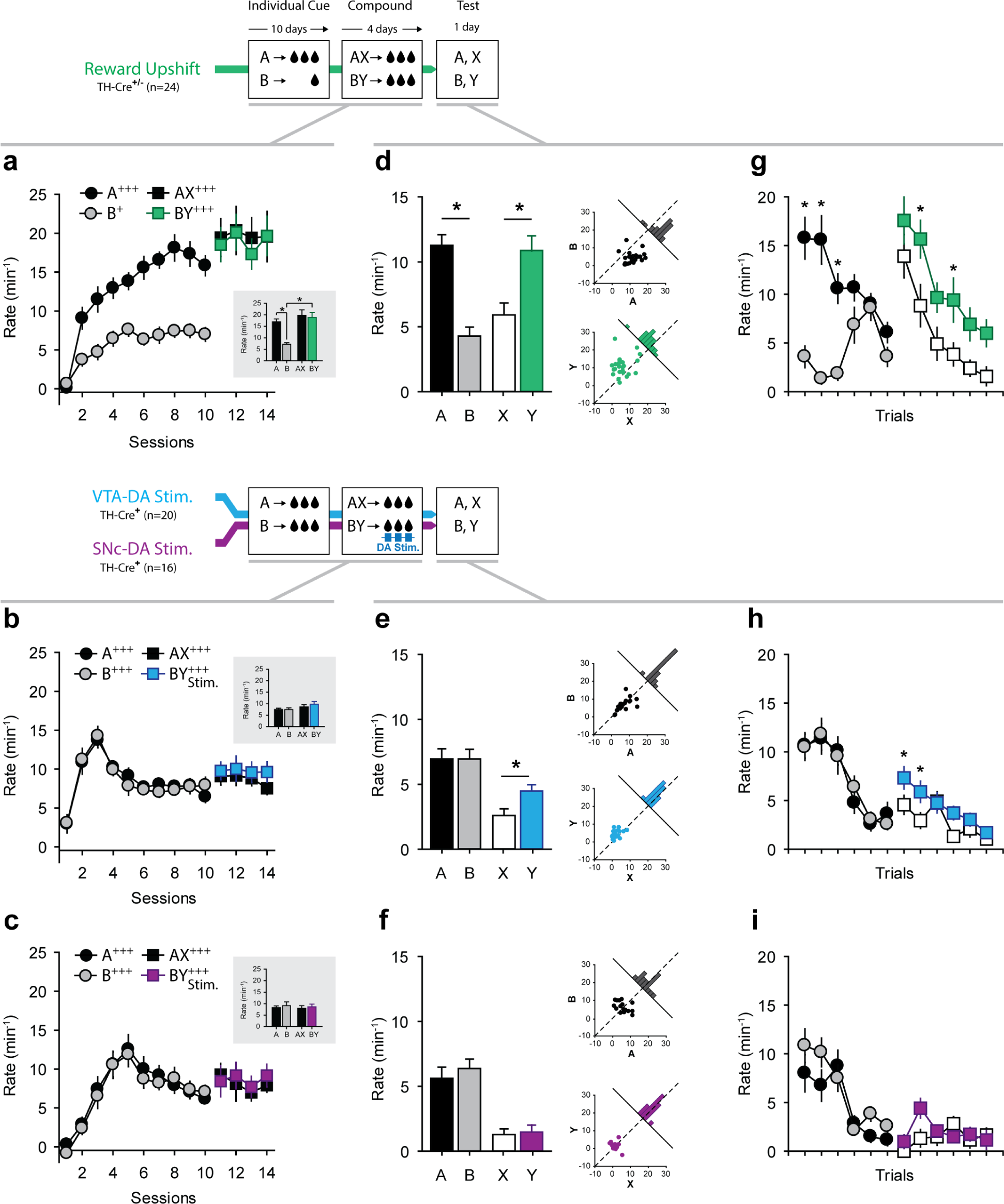
Performance in the Blocking/Unblocking task, as measured by the rate of port entries. **(a-c)** Rate of port entries during cue presentation over the course of 10 days of individual cue conditioning and 4 days of compound cue conditioning, for the reward upshift **(a)**, VTA-DA stimulation **(b)**, and SNc-DA stimulation **(c)** groups. The reported values consider only the first 9s of the cues (prior to sucrose delivery) in order to avoid contamination with the sucrose consumption period. The inserts represent the average performance over the last 4 days of individual cue conditioning (cue A and B) and the 4 days of compound cue conditioning (compound AX and BY). In the reward upshift group, responding to cue A was higher than to cue B (RM-ANOVA F_3,69_ = 17.141, P<0.001; *post hoc* Bonferroni: A vs B: T = 4.947, P<0.001), which is consistent with the magnitude of reward associated with these two cues. This difference disappeared during the compound phase as both AX and BY compounds signaled a large reward (*post hoc* Bonferroni: AX vs BY: T = 0.400, P = 1.000). No difference in the rate of port entries during the different cues was observed for the other two groups (all Ps > 0.842). **(d-f)** Rate of port entries during the final probe test, for the reward upshift **(d)**, VTA-DA stimulation **(e)**, and SNc-DA stimulation **(f)** groups. Conditioned behavior was measured during the entire 30s cue. Scatterplot inserts on the right represent individual data distribution for responses to cue A and B (top inserts), and X and Y (bottom insert). Histograms along the diagonal line represent the frequency distribution (subject counts) for the difference score in responding (A-B, or X-Y); off-centered distributions reveal higher responding to one of the cues. A two-way mixed ANOVA (Group × Cue) revealed a main effect of Group (F_2,57_ = 478.943, P<0.001) and Cue (F_3,171_ = 18.763, P<0.001) and a significant interaction between these factors (F_6,171_ = 11.929, P<0.001). Subsequent oneway RM-ANOVAs, separately conducted on each experimental group, confirmed a main effect of Cue in each group (all Ps<0.001). Post-hoc comparisons showed higher responding to Y than to X after reward upshift (T = 4.065, P<0.001), or VTA-DA stimulation (T = 2.752, P=0.048), but not after SNc-DA stimulation (T = 0.195, P = 1.000). **(g-i)** Trial-by-trial performance in the reward upshift group **(g)**, VTA-DA stimulation group **(h)**, and SNc-DA stimulation group **(i)**. A 3-way mixed ANOVA (Group × Cue × Trial) analyzed the evolution of responding over the course of the session and found an interaction between all factors (F_30,855_ = 4.605, P<0.001, after Greenhouse-Geisser correction). Results of planned contrast analyses are shown on the graphs. *P<0.05 (A *vs*. B, or X *vs*. Y; Bonferroni t-tests). Error bars = s.e.m.

**Figure S2.**
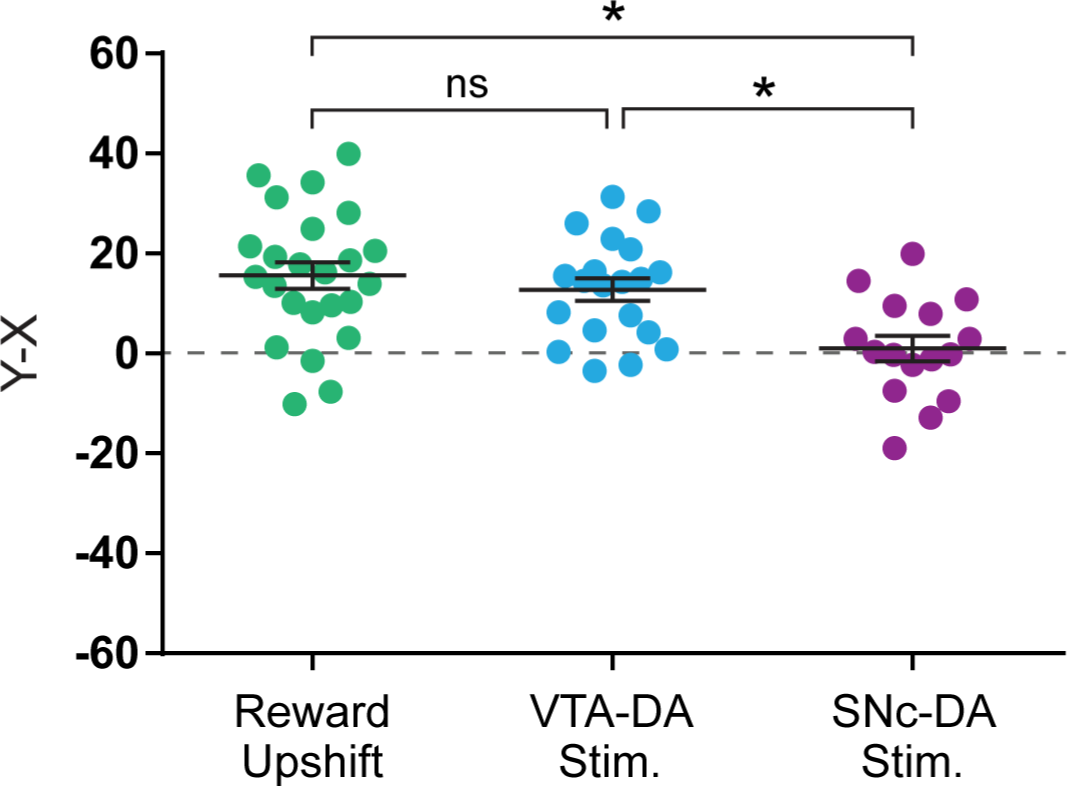
Unblocking score. To measure and compare the efficacy of the different manipulations (reward upshift, VTA- or SNc-DA stimulation), an unblocking score was calculated for all subjects. This score is defined as the difference in responding between the unblocked cue and the blocked cue (Y-X; using time in port as the measure of responding). An ANOVA conducted on the unblocking score found a significant effect of Group (F_2,57_ = 8.247, P < 0.001). *Post-hoc* tests revealed that Reward Upshift and VTA-DA stimulation resulted in comparable unblocking (Upshift *vs*. VTA-DA Stim: T = 0.817, P = 1.000), while the SNc-DA group stood different from the other two groups (SNc-DA *vs*. Upshift: T = 3.947, P < 0.001; SNc-DA vs. VTA-DA: T = 3.060, P = 0.010).

**Figure S3.**
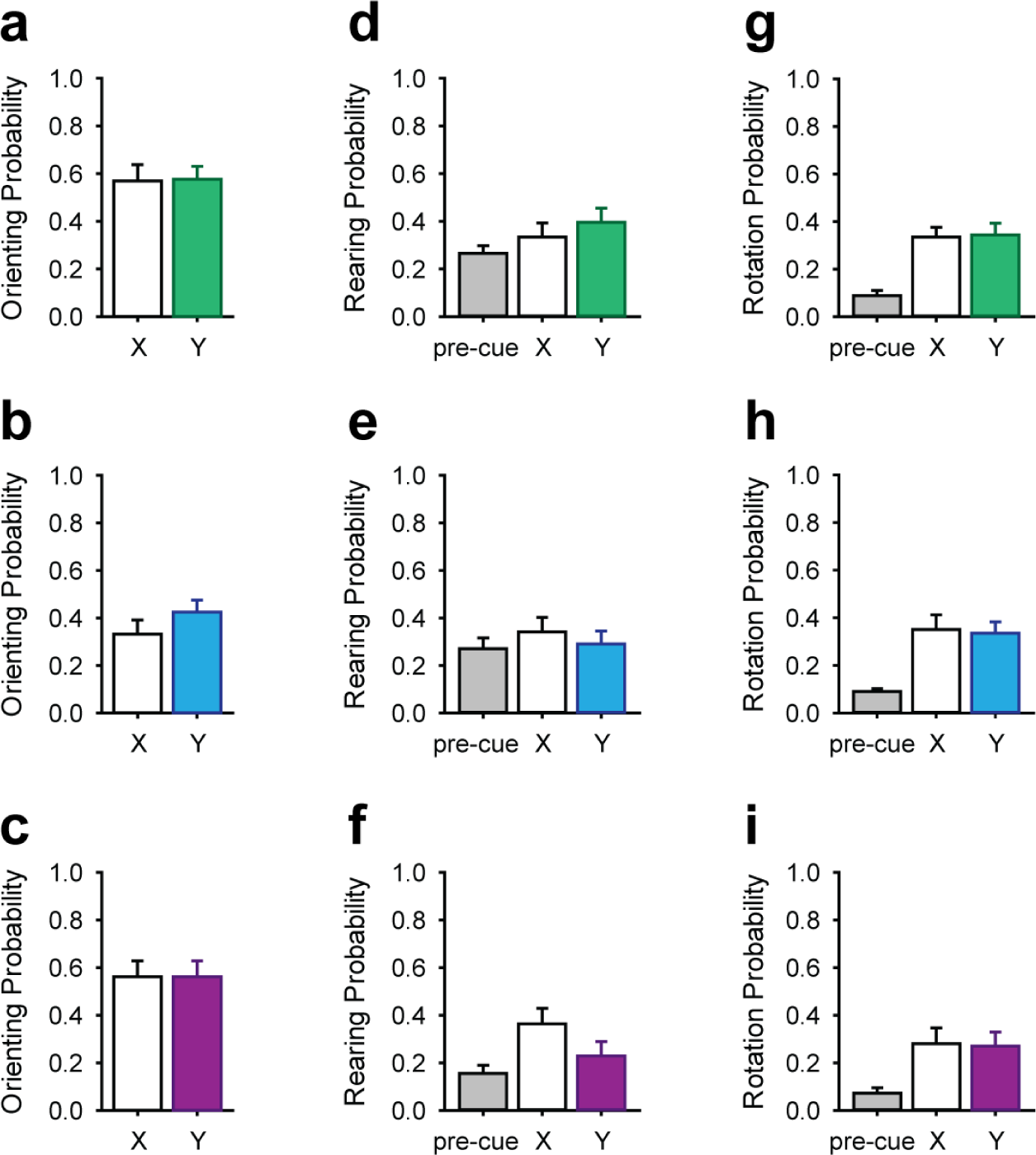
Accessory behaviors (not directed towards the reward port) evoked by the target cues X and Y during the probe test. **(a-c):** Probability of cue-evoked orienting response in the reward upshift group **(a)**, VTA-DA stimulation group **(b)**, and SNc-DA stimulation group **(c)**. A two-way mixed ANOVA (Group × Cue) conducted on the probability of cue-evoked orienting response revealed a main effect of Group (F_2,57_ = 9.646, P<0.001), but no main effect (F_1,57_ = 0.305, P = 0.583) or interaction with Cue (F_2,57_ = 0.249, P = 0.781). Post-hoc comparisons indicated that VTA-DA group displayed fewer orienting responses compared to the other two groups (VTA-DA *vs*. Upshift: T = 4.055, P<0.001; VTA *vs*. SNc: T = 3.464, P = 0.003; Upshift vs. SNc-DA: T = 0.204, P = 1.000, Bonferroni t-tests). **(d-f):** Probability of rearing in baseline (pre-cue period) and during cue presentation in the reward upshift group **(d)**, VTA-DA stimulation group **(e)**, and SNc-DA stimulation group **(f)**. A two-way mixed ANOVA (Group × Cue) conducted on the probability of rearing revealed a small but significant effect of cue presentation (F_2,114_ = 4.264, P = 0.027 after Greenhouse-Geisser correction), but no main effect (F_2,57_ = 1.224, P = 0.302) or interaction with Group (F_4,114_ = 1.328, P = 0.264). **(g-i)**: Probability of rotation in baseline (pre-cue period) and during cue presentation in the reward upshift group **(g)**, VTA-DA stimulation group **(h)**, and SNc-DA stimulation group **(i)**. A two-way mixed ANOVA (Group × Cue) conducted on the probability of rearing revealed a main effect of cue presentation (F_2,114_ = 49.060, P<0.001 after Greenhouse-Geisser correction), but no main effect (F_2,57_ = 1.781, P = 0.178) or interaction with Group (F_4,114_ = 0.472, P = 0.711, after Greenhouse-Geisser correction). Post-hoc comparisons revealed that, compared to the pre-cue period, both cues X and Y increased rotations to a similar extent (Pre-cue *vs*. X: T = 9.024, P < 0.001; Pre-cue *vs*. Y: T = 8.897, P < 0.001; X *vs*. Y: T = 0.127, P = 1.000). Error bars = s.e.m.

**Figure S4.**
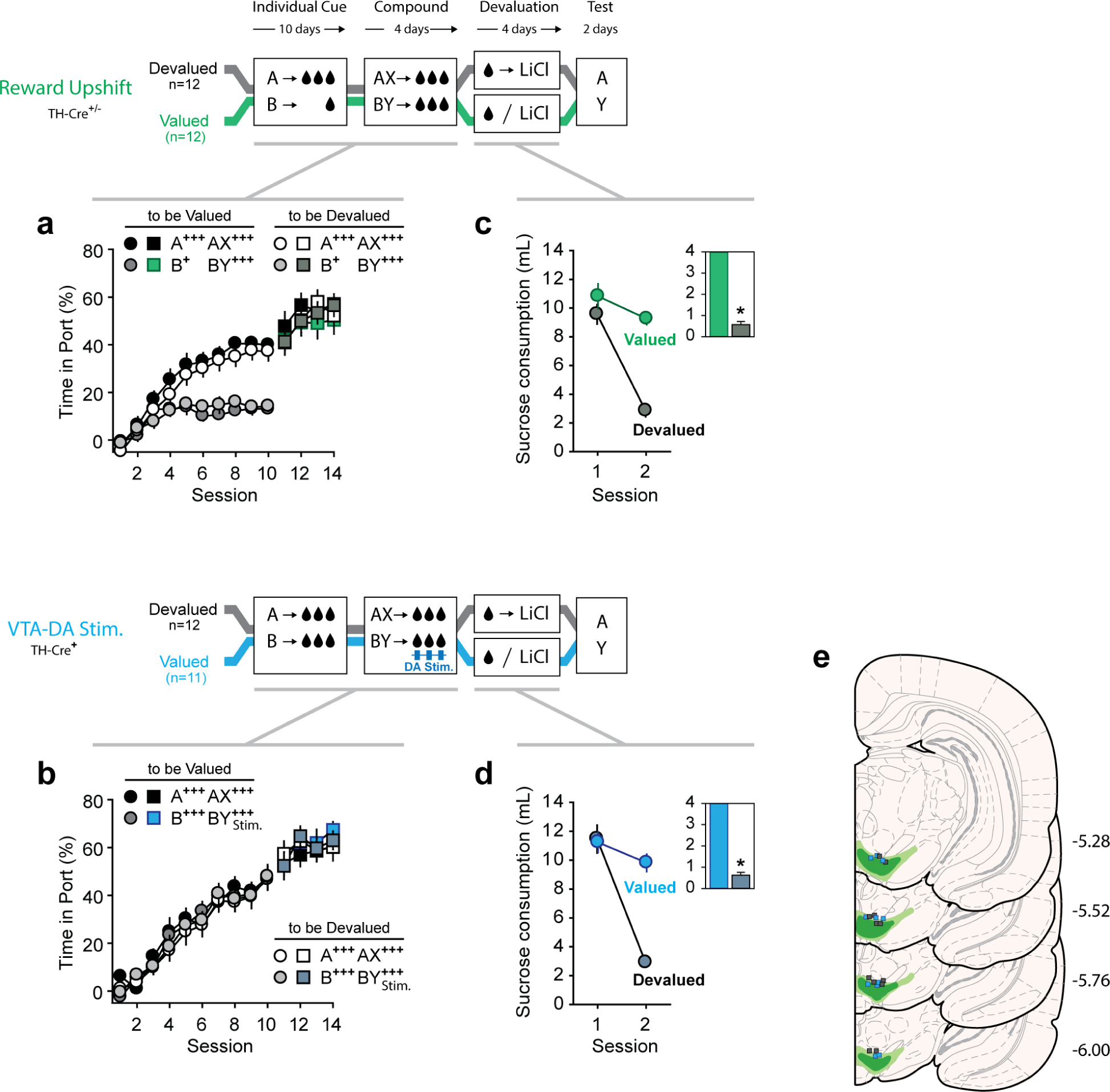
Devaluation task design. Subjects were initially trained in the unblocking task as previously described, where learning about the target cue Y was unblocked by reward upshift (top graphs) or by photoactivation of VTA-DA neurons (bottom graphs). **(a-b)**: Performance during the individual and compound cue phases (measured as time in port during cue presentation) for the Reward Upshift group **(a)** and the VTA-DA stimulation group **(b)**. At the end of the compound conditioning phase and before the final probe test, half of the rats in each group had the sucrose outcome devalued by pairing it with lithium chloride (LiCl) (devalued condition). The other half of the rats (valued condition) received similar exposure to sucrose and LiCl but on alternate days. **(c-d)** Sucrose consumption during the devaluation procedure, for the Reward Upshift group **(c)** and the VTA-DA stimulation group **(d)**. Pairing sucrose consumption with LiCl significantly reduced consumption. A 3-way mixed ANOVA (Group × Session × Devaluation) conducted on homecage sucrose consumption revealed a significant Session × Devaluation interaction (F_1,43_ = 64.384, P < 0.001) but no main effect (F1,43 = 3.406, P = 0.072) or interaction with Group (Group × Session: F_1,43_ = 1.502, P = 0.227; Group × Devaluation: F_1,43_ = 0.050, P = 0.824; Group × Session × Devaluation: F_1,43_ = 0.707, P = 0.405). Post-hoc comparisons revealed that differences between the valued and devalued condition emerged after the first injection of LiCl (day 2; valued *vs*. devalued T = 10.428, P < 0.001). The aversion for sucrose easily transferred across context and was observed during brief consumption test in the conditioning chambers (inserts). **(e)** Histological reconstruction of ChR2-YFP expression and fiber placement in the VTA. Light and dark shading indicate maximal and minimal spread of ChR2-YFP respectively. Square symbols mark the ventral extremity of the fiber implants (blue: valued group; gray: devalued group). *: P<0.05 (Devalued *vs*. Valued; Mann-Whitney Rank Sum Test). Error bars = s.e.m.

**Figure S5.**
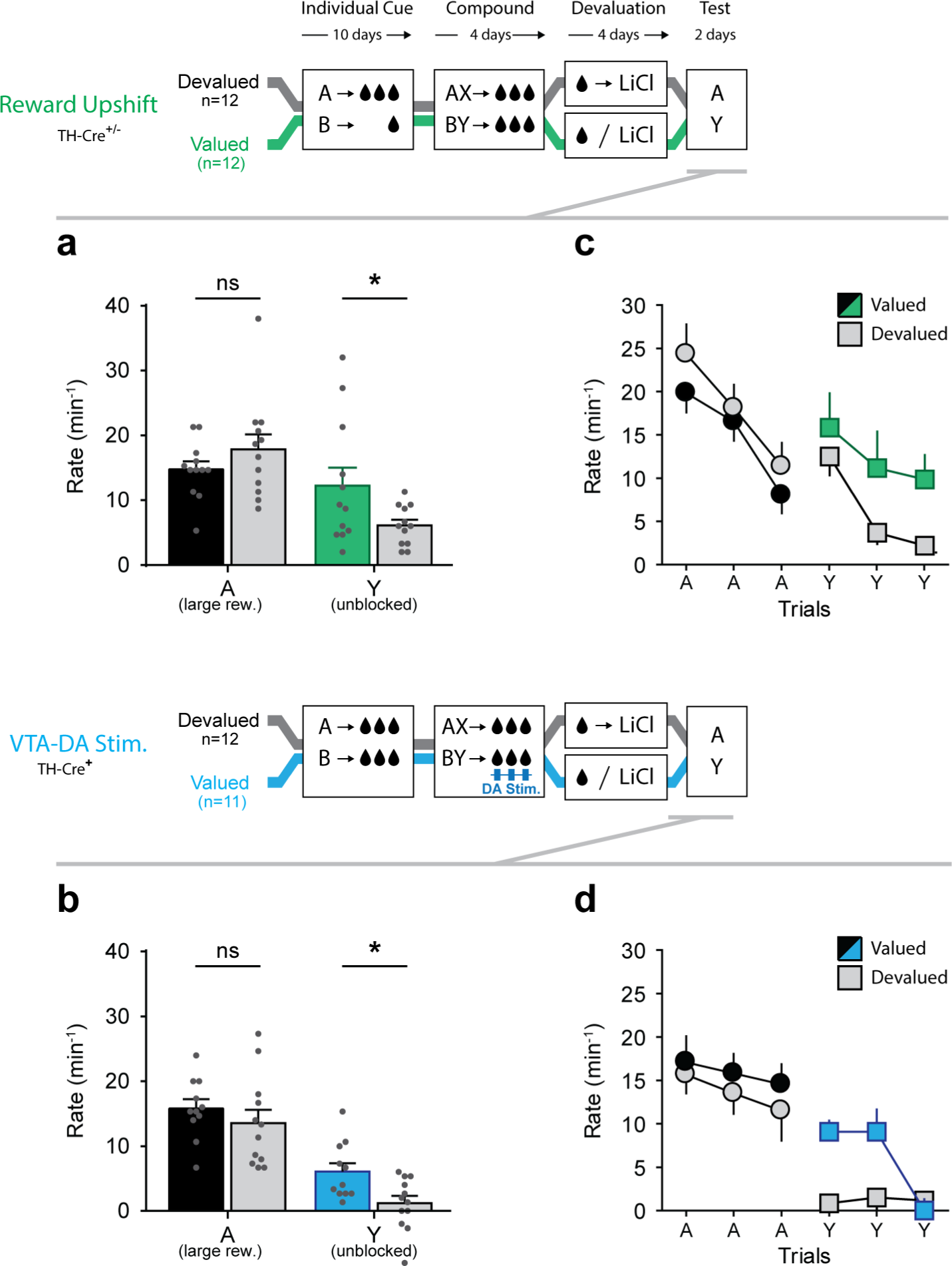
Effect of sucrose devaluation on conditioned responding during probe test, as measured by the rate of port entries. **(a, b)** Rate of port entries during cue presentation in the Reward Upshift group **(a)** and VTA-DA stimulation group **(b)**. A 3-way mixed ANOVA (Group × Devaluation × Cue) conducted on the rate of port entries during the cues revealed a main effect of Cue (F_1,43_ = 43.631, P < 0.001), Devaluation (F_1,43_ = 5.236, P = 0.027), and significant interaction between these two factors (F_1,43_ = 4.679, P < 0.036). This interaction was due to the significant reduction of responding to cue Y (T = 3.046, P = 0.003), but not to cue A (T = 0.248, P = 0.805) following sucrose devaluation. The ANOVA also revealed a main effect of Group (F_1,43_ = 10.588, P = 0.002) as responding was overall higher in the Reward Upshift group; however no interaction with Group was found significant (Group × Cue: F_1,43_ = 2.018, P = 0.163; Group × Cue × Devaluation: F_1,43_ = 1.464, P = 0.233). Planned contrast analysis independently confirmed that, in both groups, sucrose devaluation reduced responding to cue Y (Reward Upshift: T= 2.102, P = 0.047; VTA-DA Stim.: T= 2.779, P = 0.011) but not to cue A (Reward Upshift: T= 1.200, P = 0.247; VTA-DA Stim.: T= 0.903, P = 0.377). **(c-d)** Trial-by-trial performance in the reward upshift group **(c)** and VTA-DA stimulation group **(g)**. *P<0.05 (Devalued *vs*. Valued). Error bars = s.e.m.

**Figure S6.**
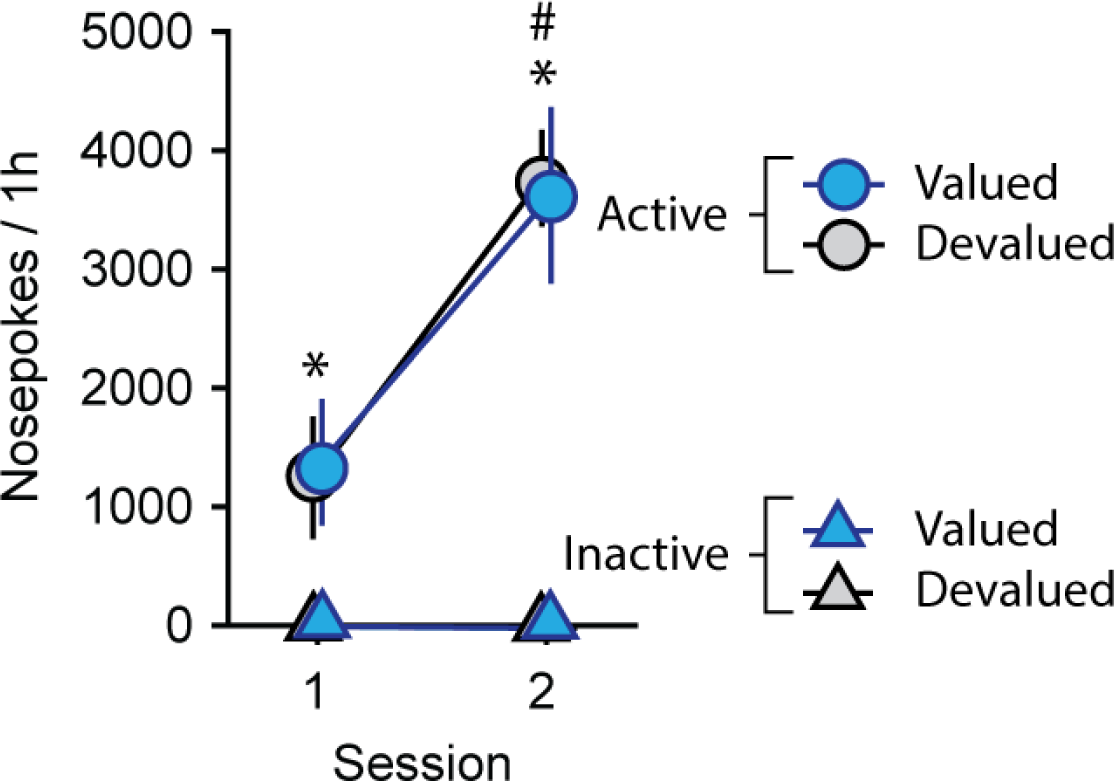
VTA-DA intracranial self-stimulation in Devalued and Valued groups. Responses at the active and inactive nosepoke over the course of two daily 1-h intracranial self-stimulation sessions. A 3-way mixed ANOVA (Devaluation × Session × Nosepoke) revealed a significant preference for the active nosepoke (main Nosepoke effect: F_1,21_ = 57.926, P<0.001) and a Nosepoke × Session interaction (F_1,21_ = 35.514, P<0.001) as responses at the active nosepoke increased over days (T = 8.527, P < 0.001, Bonferroni t-test) while responding at the inactive nosepoke remained virtually absent (T = 0.0495, P = 0.961, Bonferroni t-test). Critically, we found no main effect (F_1,21_ = 0.000, P = 0.986) or interaction with Group (F_1,21_ < 0.142, Ps > 0.710) indicating that VTA-DA stimulation is equally reinforcing in the Valued and Devalued groups. *P < 0.05, Active vs. Inactive Nosepoke; ^#^P < 0.05, Session 1 vs. Session 2 (active nosepoke). Error bar and error bands = s.e.m.

